# Bearskin2 mediates the coordinated secretion of xylogalacturonan and root cap polygalacturonase in Arabidopsis border-like cells

**DOI:** 10.1101/2023.05.21.541628

**Authors:** Zhongyuan Liu, Pengfei Wang, Tatsuaki Goh, Keiji Nakajima, Byung-Ho Kang

**Affiliations:** School of Life Sciences, Centre for Cell and Developmental Biology and State Key Laboratory of Agrobiotechnology, The Chinese University of Hong Kong, Shatin, New Territories, Hong Kong, China; Graduate School of Science and Technology, Nara Institute of Science and Technology, 8916-5 Takayama, Ikoma, Nara 630-0192, Japan

## Abstract

Border-like cells (BLCs) are sheets of cells that are continuously sloughed off and replenished at the Arabidopsis root cap surface. *ROOT CAP POLYGALACTURONASE (RCPG)* encodes a putative pectinase involved in BLC shedding. Xylogalacturonan (XGA) is a pectic polysaccharide whose synthesis is associated with cell detachment and secreted separately from other cell wall polysaccharides. *BEARSKIN1 (BRN1)* and *BRN2* are *Arabidopsis* NAC family transcription factors, and *RCPG* expression is inhibited in *brn1/2*. To explore the link between XGA and RCPG, we examined XGA synthesis in *Arabidopsis* lines with altered RCPG levels. We found that RCPG was contained in XGA-carrying vesicles budding from the *trans*-Golgi, but XGA synthesis was not affected in the *rcpg* mutant. XGA was absent in BLCs of *brn2*, but not of *brn1*, indicating that *BRN2* is necessary for XGA synthesis. Overexpression of functional RCPG-GFP (*oeRCPG-GFP*) caused upregulation of *BRN2*, ectopic XGA synthesis, overaccumulation of endogenous RCPG, and accelerated BLC turnover, suggesting a positive regulatory loop between RCPG and BRN2. Inactivation of *BRN2* in *oeRCPG-GFP* suppressed RCPG-GFP expression, excess RCPG, and XGA synthesis. Our data provide evidence that XGA and RCPG are secreted together and that BRN2 controls XGA synthesis, which facilitates RCPG export and BLC separation.

## Introduction

The root cap covers the tip of the plant root, protecting the root meristem cells, directing root growth, and secreting mucilage into the soil. The root cap is under a constant cell flux, new root cap cells arise in the basal root meristem, differentiate, and replace the root cap surface cells that are sloughed off (Barlow, 2002; Sievers et al., 2002; Arnaud et al., 2010). Border cells are root cap surface cell individually detached from the root cap (Hawes and Lin, 1990; Driouich et al., 2007). In Arabidopsis, border cells constitute a cell layer that is shed together, and they are called border-like cells (BLCs) (Vicre et al., 2005; Durand et al., 2009; Driouich et al., 2010). Border cells and BLCs lie at the forefront of root tip and secrete molecules to amend the rhizosphere conducive to root growth (Iijima et al., 2004; Driouich et al., 2013; Maeda et al., 2019).

The root cap exhibits high secretory activity, as evidenced by the presence of hypertrophied Golgi stacks (Whaley et al., 1959; Spink and Wilson, 1968; Staehelin et al., 1990). The mucilage of the root cap primarily comprises pectin polysaccharides and proteoglycans (Read et al., 1999; Maeda et al., 2019; Castilleux et al., 2020; Ropitaux et al., 2020). Golgi stacks in border cells/BLCs and peripheral cells, precursors of border cells/BLCs have swollen cisternae where synthesis of pectin and addition of polysaccharide moieties of proteoglycans occur (Wang et al., 2017; Wang and Kang, 2018). The unique architecture of the Golgi stack in border cells and its assembly process have piqued the interest of plant cell biologists. Recently it was shown that cell maturation and programmed cell death in the root cap involve reorganization of the cytoplasm and activation of autophagy, recapitulating the drastic intracellular changes along the developmental gradient in the root cap (Feng et al., 2022; Goh et al., 2022).

The Arabidopsis root cap consists of two distinct regions: the central columella root cap (CRC) and the lateral root cap (LRC) that surrounds it (Dolan et al., 1993; Barlow, 2002). The detachment of BLCs in these regions is mediated by separate mechanisms. In the LRC, BLC shedding is triggered by the activation of programmed cell death (PCD), which depends on a NAC-type transcription factor, SOMBRERO (SMB) (Bennett et al., 2010; Fendrych et al., 2014). In the CRC, BLC separation involves the activity of a putative pectin-digesting enzyme, ROOT CAP POLYGALACTURONASE (RCPG). Two NAC-type transcription factors, BEARSKIN 1 (BRN1) and BRN2 exhibit high sequence similarity and they are specifically expressed in the outer cell layers of the root cap. They contribute to BLC shedding from CRC via activating RCPG, and it has been demonstrated that BRN1 binds to the promoter region of *RCPG* (Kamiya et al., 2016).

Xylogalacturonan (XGA) is a type of pectic polysaccharide that features a homogalacturonan backbone composed of (α-(1→4)-linked d-galacturonic acid), with a β-d-xylose substitution occurring at the O-3 position (Zandleven et al., 2006). XGA has been found in various types of plant cells, but its epitope is particularly abundant in the root cap and seed coat, which are regions where cell walls break down and cells are shed (Willats et al., 2004; Zandleven et al., 2007; Wang et al., 2017; Wang and Kang, 2018). XGA is secreted from border cells/BLCs of alfalfa, pea, maize, and Arabidopsis. Our electron microscopy analysis of border cells/BLCs indicate that XGA accumulates in the swollen margins of *trans*-Golgi cisternae where it is sorted into vesicles instead of transported to the *trans*-Golgi network (TGN) (Wang et al., 2017; Wang and Kang, 2018). Given that pectin polysaccharides are crucial for the construction and maintenance of the primary cell wall and are abundant in middle lamellar that is responsible for plant cell adhesion, XGA secretion from BLCs that are detached from the root cap is intriguing (Bouton et al., 2002; Mouille et al., 2007; Durand et al., 2009; Du et al., 2022).

Based on the BLC-specific expression of RCPG and the secretion of XGA, we hypothesized that there is a coordinated regulation between RCPG and the machinery responsible for XGA synthesis. To test this hypothesis, we examined the expression patterns and subcellular localizations of RCPG and XGA in Arabidopsis lines exhibiting abnormal *RCPG* expression. Our findings indicated that RCPG is a protein cargo of vesicles carrying XGA, and *BRN2,* not *BRN1,* is required for XGA synthesis. Furthermore, we observed that overexpression of RCPG-GFP led to ectopic XGA synthesis and excess RCPG accumulation, which were dependent on *BRN2*. These results suggest that *BRN2* couples *RCPG* expression and XGA synthesis in the Arabidopsis BLC.

## Results

### Expression of *RCPG* is linked to XGA secretion in the *Arabidopsis* root cap

At six days after germination (DAG), BLC release and XGA secretion were clearly detected in the Arabidopsis root cap. However, no BLCs or XGA were observed in the young root cap at 2 DAG (Fig, 1 A-B). A transgenic line expressing an RCPG-RFP fusion protein with the RCPG native promoter *(pRCPG:RCPG-RFP)* did not exhibit RFP fluorescence at 2 DAG. By contrast, RCPG-RFP accumulated in BLCs at 6 DAG, indicating an association between RCPG and BLC (Figure 1 C-D). In BLCs of 6 DAG root caps, hypertrophied margins of trans-Golgi carrying XGA were seen in Golgi stacks, but not in those of 2 DAG root cap cells (Figure 1 E-H). Immunoblot analysis using an RCPG antibody detected RCPG polypeptides in protein extracts from 6 DAG root cap samples, but not in 2 DAG root cap samples (Figure 1 I-J), further supporting the association between RCPG expression and XGA secretion.

**Figure 1.**
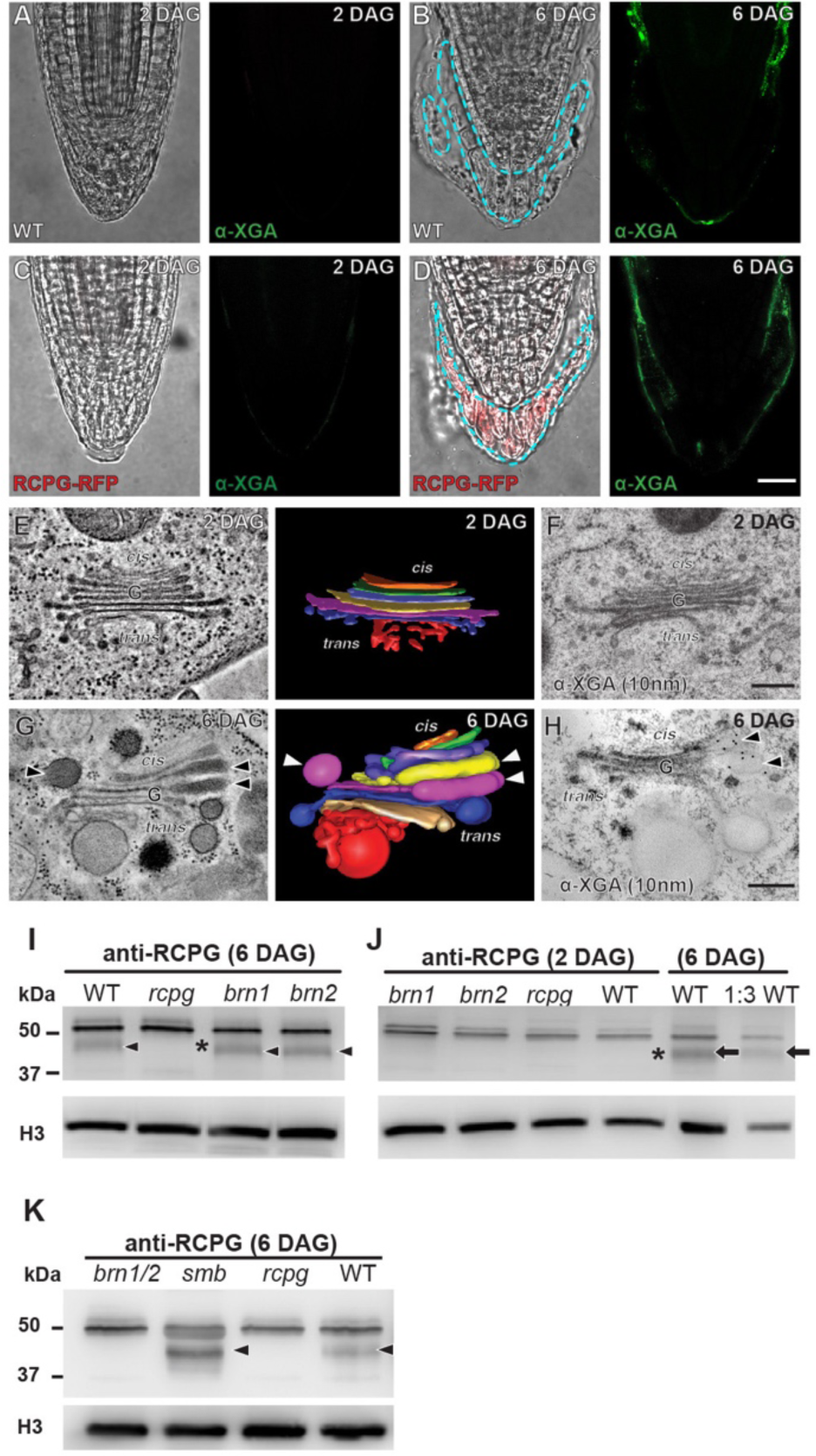
Expression of RCPG and XGA synthesis in Arabidopsis BLCs. **A-D**, Whole mount immunofluorescence with a XGA antibody of wild-type (WT; A and B) and *pRCPG:RCPG-RFP* transgenic line (C and D) at 2 and 6 DAG. BLCs separating from the root cap surface are marked with blue dashed lines in B and D. Root caps do not shed cells and secrete XGA at 2 DAG (A and C). Bar = 25 μm. **E-H**, Electron tomography and immunogold labeling of Golgi stacks in root cap surface cells at 2 DAG (E-F) and in BLCs at 6 DAG (G-H). Golgi stacks in BLCs at 6 DAG have swollen cisternal margins containing XGA (arrowheads in G and H). Bars = 200 nm. **I-J**. Immunoblot analysis of root cap protein extracts at 6 DAG (I) and 2 DAG (J) with a RCPG antibody. Arrowheads mark RCPG polypeptides. The RCPG polypeptide is absent in the protein samples from *rcpg* root caps at 6 DAG (asterisk in I). For the 2 DAG immunoblot, two dilutions of protein samples from 6 DAG WT root cap were analyzed as the positive control (arrows). **K.** Immunoblot analysis of protein extracts from *brn1/2*, *smb*, and *rcpg* mutant root cap samples. *brn1* and *brn2* single mutant samples have RCPG at but *brn1/2* double mutant does not at 6 DAG. Arrowheads mark RCPG polypeptides.

### RCPG is a protein cargo of XGA-carrying vesicles

We investigated the localization of RCPG in *pRCPG:RCPG-RFP* root cap cells using immunogold labeling. To simplify RCPG localization, we created an Arabidopsis line, *oeRCPG-GFP*, in which RCPG-GFP (C-terminal fusion) was overexpressed with the *ubiquitin 10* (*UBQ10*) promoter (Fig. 2A). This was necessary because the RCPG native promoter exhibited fluctuations in its activity, which complicated RCPG localization (Fig. 3). The inhibited BLC separation from the *rcpg* root cap was rescued in by transformation with *oeRCPG-GFP* (Fig. S1H). When we carried out immunogold labeling in the two transgenic lines, RFP and GFP-specific gold particles were associated with XGA-carrying vesicles budding from the trans-Golgi (Fig. 2 B-C). However, no RFP or GFP-specific gold particles associated with the trans-face of the Golgi where the trans-Golgi network (TGN) arises.

**Figure 2.**
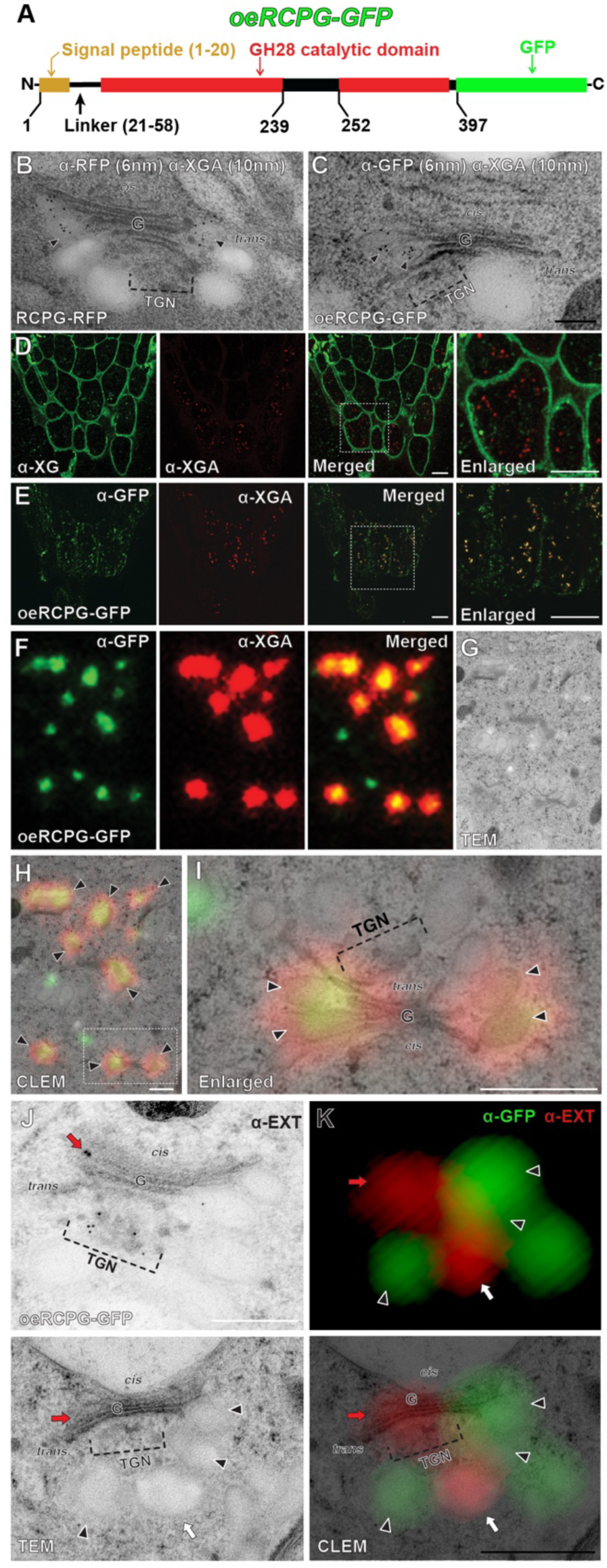
RCPG is a protein cargo of XGA-carrying vesicles. **A**. domain architecture of RCPG-GFP fusion protein in the *oeRCPG-GFP* line. RCPG is 397 amino acid long, consisting of a signal peptide (yellow), a linker domain (black line), and a GH28/catalytic domain (red). The active site in the GH28 domain is marked (black box). **B-C**. double immunogold labelling with RFP/GFP and XGA antibodies in *pRCPG:RCPG-RFP* (B) and *oeRCPG-GFP* (C). RCPG-RFP and RCPG-GFP colocalized with XGA in vesicles budding from the *trans*-Golgi (arrowheads). Bars = 200 nm. **D.** double immunofluorescence of XGA and xyloglucan (XG). Bars = 50 μm. **E.** double immunofluorescence of XGA and GFP in the root cap of *oeRCPG-GFP*. XGA-specific fluorescence (α-XGA) overlapped with anti-GFP but not with XG. Bars = 50 μm. **F-H.** CLEM localization of RCPG-GFP and XGA. GFP and XGA-specific punctate spots overlap (F). The fluorescence puncta correspond to swollen vesicles of the *trans*-Golgi (H) when fluorescence (H) and electron (G) micrographs were aligned. **I.** enlarged image of the boxed area in H. Arrowheads in H and I indicate RCPG-GFP colocalizing with XGA. Bars in H and I = 500 nm. J. Immunogold labeling of extensin. Gold particles are seen in Golgi cisternae (red arrow) and in trans-Golgi network (TGN) compartments. K. CLEM localization of RCPG-GFP (green) and extensin (red). Arrowheads, white arrow, and red arrow denote trans-Golgi vesicles, TGN vesicles, and Golgi cisternae, respectively and they point to same locations in each micrograph. Extensin-specific puncta overlap with Golgi cisternae, TGN, and TGN derived vesicles, distinct from XGA vesicles. Scale bars in J and K: 500 nm.

**Figure 3.**
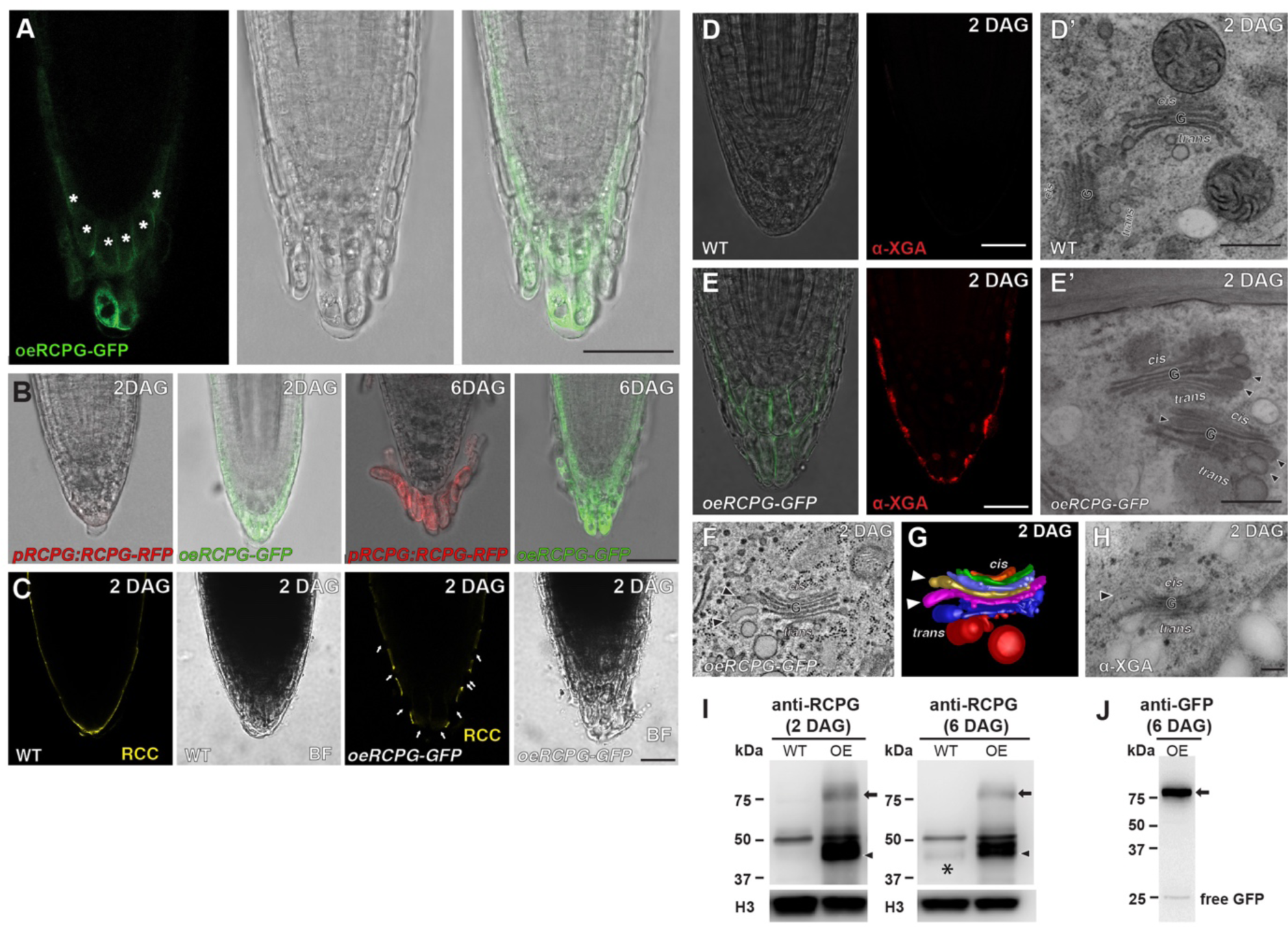
Ectopic XGA synthesis and excess *RCPG* accumulation in *oeRCPG-GFP*. **A.** RCPG-GFP is detected in the cell layer inside BLCs (asterisks) in the RCPG-GFP overexpressor line (*oeRCPG-GFP)*. Bar = 50 μm. **B.** Expression of RCPG-GFP regulated by the native *RCPG* promoter (*pRCPG:RCPG-RFP*) and by the *Ubq10* promoter (*oeRCPG-GFP)* at 2 DAG and 6 DAG. RCPG-GFP is observed in 2 DAG root cap of *oeRCPG-GFP*. Bars = 50 μm. **C**. Root cap cuticle (RCC) of wild-type (WT) and *oeRCPG-GFP* at 2 DAG. Arrowheads indicate cracks in RCC at the *oeRCPG-GFP* root cap surface. Bar = 25 μm. **D-E**. Whole mount immunofluorescence and TEM micrographs of wild-type (WT; D) and *oeRCPG-GFP* (E) at 2 DAG. Arrowheads indicate XGA-carrying vesicles budding from the *trans*-Golgi in *oeRCPG-GFP* root cap surface cell (B’). Bars in A and B = 25 μm. Bars in electron micrographs = 500 nm. **F-H.** Electron tomographic slice (F) and 3D model (G) images of a Golgi stack in a root cap surface cell of *oeRCPG-GFP* at 2 DAG. **E.** Immuno-electron micrograph of a root cap surface cell in *oeRCPG-GFP* at 2 DAG with a XGA antibody. Arrowheads denote *trans*-Golgi vesicles containing XGA. Bar = 200 nm. **F-G**, Immunoblot analysis of WT and *oeRCPG-GFP* (OE) with a RCPG (I) and a GFP antibody (J). Asterisk marks endogenous RCPG in WT. Arrows and arrowheads indicate RCPG-GFP and endogenous RCPG in *oeRCPG-GFP* (OE) lanes, respectively. Histone H3 amounts were assessed for loading control.

We performed correlative light and electron microscopy (CLEM) analysis to confirm the localization of RCPG-GFP in XGA-carrying vesicles. First, we collected sections from *oeRCPG-GFP* sample blocks prepared for transmission electron microscopy (TEM) and processed them for immunofluorescence microscopy using anti-XGA and anti-GFP antibodies. After imaging with fluorescence microscopy, we post-stained the sections and captured electron micrographs of the regions where XGA or GFP fluorescence was detected. Our observations revealed that GFP fluorescence overlapped with XGA-specific fluorescence in BLCs (Fig. 2E). When fluorescence micrographs and electron micrographs were aligned, puncta positive for both XGA and GFP corresponded to swollen vesicles budding from the trans-Golgi (Fig. 2 F-I). Notably, xyloglucan (XG), which localizes to the trans-Golgi network (TGN), did not colocalize with XGA (Fig. 2D). Extensin is a cell wall glycoprotein that promotes cell wall expansion. Double immunofluorescence microscopy of extensin and RCPG-GFP showed that its fluorescence did not overlap with RCPG-GFP in the Golgi (Fig. 2J). CLEM and immunogold labeling indicated that extensin was associated with Golgi cisternae, TGN, and TGN-derived vesicles, revealing that extensin and RCPG-GFP are delivered to the cell wall through distinct vesicles (Fig. 2 J-I). These results support the conclusion that RCPG-GFP is packaged into XGA-carrying vesicles.

To investigate how RCPG is sorted into XGA vesicles, we generated Arabidopsis plants expressing truncated RCPG-GFPs under the *Ub10* promoter. One form lacked the catalytic domain, RCPG(ΔC)-GFP, while the other lacked the linker region between the signal peptide and catalytic domain, RCPG(ΔL)-GFP (Fig. 2A). The full-length RCPG-GFP was observed in root cap cells, particularly in BLCs and their immediate precursor cells but the truncated RCPG-GFPs accumulated in non-root cap cells (Fig. S1 A-C). While the full-length RCPG-GFP was secreted into the cell wall, the truncated RCPG-GFPs were retained in the cytoplasm (Fig. 1S E-G). Fluorescence microscopy and immunogold labeling of the truncated RCPG-GFPs revealed that RCPG(ΔC)-GFP is associated with the cis-Golgi, and RCPG(ΔL)-GFP is associated with the ER (Fig. S2). However, the abnormal forms of RCPG-GFP failed to rescue the impaired BLC detachment from the rcpg root cap (Fig. S1H).

### Overexpression of RCPG-GFP induced ectopic XGA synthesis and excess accumulation of endogenous RCPG

The *oeRCPG-GFP* line showed GFP fluorescence in peripheral cells inside the border-like cells (BLCs), as indicated by asterisks in Fig. 3A, in contrast to the pRCPG:RCPG-GFP of which fluorescence was confined in BLCs (Fig. 3B). Additionally, GFP fluorescence was observed in root cap surface cells at 2 days after germination (DAG), when the native RCPG promoter is inactive (Fig. 3B). In young roots, the cuticle covers the Arabidopsis root cap before BLC shedding begins (Berhin et al., 2019). The overexpression of RCPG-GFP disrupted the cuticle layer of 2 DAG root cap, indicating an alteration in the structure of the cuticle and outer cell wall due to ectopic RCPG-GFP (Fig 3C). Despite that RCPG-GFP expression was driven by the *Ub10* promoter, RCPG-GFP was restricted to the root cap of *oeRCPG-GFP* at 2 and 6 DAG (Fig. 3 A-B).

As RCPG-GFP was produced in the 2 DAG root cap of the oeRCPG-GFP line, we investigated whether early expression of RCPG-GFP could activate XGA synthesis. Immunofluorescence labeling showed that XGA epitopes accumulated on the root cap surface of the overexpression line at 2 DAG (Fig. 3 D-E). In electron micrographs/electron tomograms of root cap cells, Golgi stacks had swollen trans-cisternae (Fig. 3 F-G) associated with XGA-specific immunogold particles (Fig. 3H). In wild-type root caps, BLCs are not shed and no XGA is synthesized at 2 DAG (Fig. 3D). Immunoblot analysis with an RCPG-specific antibody indicated that endogenous RCPG is synthesized in 6 DAG but not in 2 DAG root tip samples (asterisk in Fig. 3I). Intriguingly, RCPG accumulated in the oeRCPG-GFP line at 2 DAG, and its level was higher than in wild-type at 6 DAG (arrowheads in Fig. 3I). The excess RCPG in the oeRCPG-GFP line was not a degradation product of RCPG-GFP, as only a small amount of free GFP was observed in the RCPG-GFP overexpression line by immunoblot analysis with a GFP antibody (Fig. 3J). Overexpression of truncated RCPG-GFPs did not lead to XGA synthesis or accumulation of RCPG at 2 DAG (Fig. S3).

### Overexpression of RCPG-GFP accelerated BLC renewal, altered the root cap cell wall morphology

To characterize the link between BLC shedding and RCPG expression, we monitored root cap of *pRCPG:RCPG-RFP* over four days with time-lapse microscopy. BLCs were lost approximately once in every 60 hours and RCPG-RFP was observed in the central BLCs from ∼20 hours before the separation (Fig. 4A, Video S1). RCPG-RFP levels displayed continued increase, peaking in separating BLCs and newly exposed BLCs did not have RCPG-RFP (Fig. 4B, dashed boxes). RCPG-GFP was constitutively expressed in the root cap of *oeRCPG-GFP* (Fig. 4C, Video S2). Interestingly, BLC turnover cycle accelerated BLC turnover (Fig. 4D, dashed boxes).

**Figure 4.**
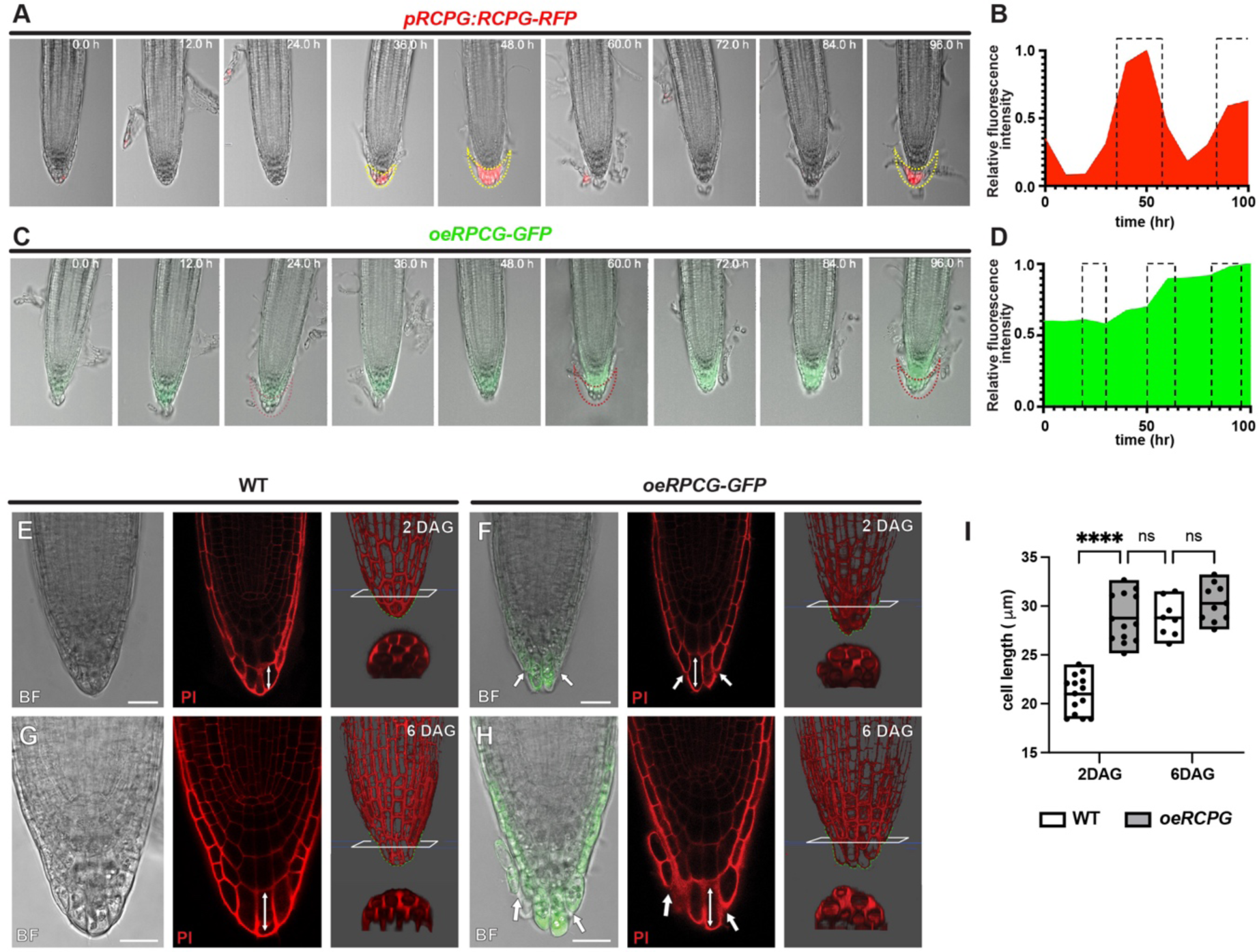
Accelerated root cap cell turnover and altered root cap morphology of *oeRCPG-GFP*. **A-D**, Time-lapse imaging of *pRCPG:RCPG-RFP* (A-B) and *oeRCPG-GFP* (C-D). 0.0 h represents the time point when the first BLC separation occurred. Dashed lines indicate BLCs being sloughed off. Relative fluorescence intensity of *pRCPG:RCPG-RFP* (B) and *oeRCPG-GFP* (D) over the imaging periods. Relative fluorescence intensity was measured by ImageJ based on Fig. 4 A and C. Dash boxes in B and D mark time windows when BLCs were released. The fluorescence intensity profile displayed cyclical dynamics in *pRCPG:RCPG-RFP* while it remained high levels throughout the 100 hrs of imaging in *oeRCPG-GFP*. **E-H**, 2 DAG and 6 DAG root tip of WT and *oeRCPG-GFP* stained by propidium iodide (PI). Double-headed arrows were added to assist comparison of lengths of BLCs under the central columellar root cap. Arrows denote depressions in cell wall profiles of the *oeRCPG-GFP* BLCs. 3D models generated from PI-stained root cap samples of WT and *oeRCPG-GFP* at 2 DAG and 6 DAG. Green dash lines delineate the root cap outlines that are irregular in *oeRCPG-GFP* in comparison to WT. In each panel, bottom images show cross-section through the root cap marked with white boxes in the upper images. Bars = 25 μm. **J**, Lengths of BLCs under the central columellar root cap in WT *and oeRCPG-GFP* at 2 DAG and 6 DAG (double-headed arrows in H-L). The BLC lengths reached 30 μm in *oeRCPG-GFP* root caps at 2 DAG, while the lengths were significantly shorter in WT (****, p<0.0001). Analysis was performed by two-way ANOVA (ns, not significant)

When the cell wall was stained with propidium iodide (PI), root cap surface cells in *oeRCPG-GFP* were elongated, and the surface cell wall had depressions at cell junctions compared to the wild-type. 3D projected views generated from image stacks clearly revealed the irregular cell wall outlines of the *oeRCPG-GFP* root tip at 2 and 6 DAG (Fig. 4 E-I).

### *BRN2*, not *BRN1,* is required for XGA synthesis in BLCs

The expression of RCPG and shedding of BLCs are impaired in the *brn1/2* double mutant root cap, as reported by Kamiya *et al*. in 2016. Immunoblot analysis using the RCPG antibody did not detect any RCPG in the *brn1/2* (Fig. 1K). Furthermore, XGA was absent in the BLCs of the *brn1/2* mutant, and their Golgi stacks did not exhibit trans-Golgi swellings (Fig. 5 A-D). In contrast, the *rcpg* mutant root cap cells displayed normal XGA synthesis. XGA-carrying vesicles were observed budding from the trans-Golgi cisternae in BLCs of *rcpg*, providing evidence that RCPG expression is not necessary for XGA synthesis (Fig. 5 E and F).

**Figure 5.**
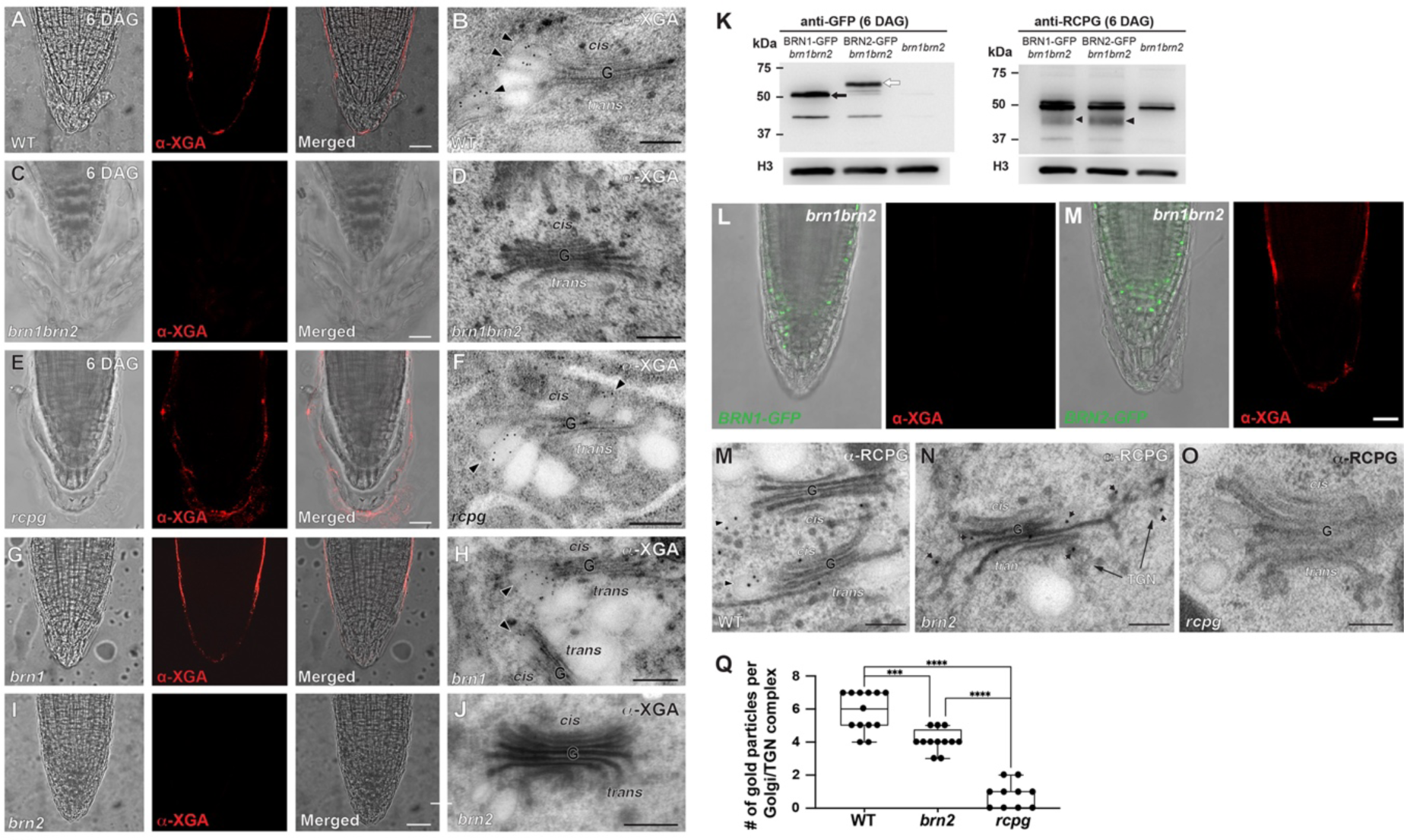
BRN2 is required for XGA synthesis and Golgi hypertrophy in BLCs. **A-J**. Whole mount immunofluorescence and immunogold labeling of Arabidopsis root cap samples (6 DAG) with a XGA antibody. XGA-specific fluorescence is missing in *brn1/2* (C) and *brn2* (I). Bars in A, C, E, G, and I (merged panels) = 25 μm. The absence of XGA on the root cap surface matched with the lack of *trans*-Golgi swelling associated with XGA-specific gold particles in D (*brn1/2*) and J (*brn2*). Bars in B, D, F, H and J (electron micrographs) = 500 nm. **K.** Inhibited *RCPG* expression in *brn1/2* rescued by expression of *BRN1-GFP* or *BRN2-GFP* with the *35S* promoter. Immunoblot with an anti-GFP antibody shows expression of BRN1-GFP (51 kDa, black arrow) and BRN2-GFP (56 kDa, white arrow). The RCPG polypeptides (arrowheads) are discerned in the *BRN1-GFP/brn1/2* and *BRN2-GFP/bran1/2* lanes, but not in the *brn1/2* lane of the immunoblot with the anti-RCPG antibody. **L-M.** XGA synthesis is restored by transforming *brn1/2* with *BRN2-GFP.* BRN2-GFP rescued the XGA synthesis defect of *brn1/2* (M) but BRN1-GFP did not (L). **N-P.** Immunogold labeling of RCPG in wild-type (WT; N), *brn2* (O), and *rcpg* (P). RCPG is associated with Golgi as well as *trans*-Golgi network (TGN) cisternae in *brn2* BLCs that do not produce XGA-carrying vesicles. Golgi stacks in *rcpg* BLCs produce XGA-carrying vesicles but lack immunogold particles, verifying the specificity of α-RCPG. **Q.** Whisker plot showing numbers of gold particles associated with Golgi/TGN complexes in WT, *brn2*, and *rcpg* BLCs. Scale bars in M-O: 200 nm

To identify which of the two *BRNs*, *BRN1* or *BRN2*, regulates XGA synthesis, we isolated *brn1* and *brn2* single mutant lines. XGA synthesis and secretion remained unaffected in the *brn1* mutant BLCs (Fig. 5 G and H). XGA synthesis was inhibited in the *brn2* BLCs, which resulted in the absence of XGA secretion and XGA vesicles in their Golgi stacks (Fig. 5 I and J). Immunoblot analysis using the RCPG antibody indicated that the levels of RCPG were not reduced in *brn1* and *brn2*, indicating that either *BRN1* or *BRN2* is sufficient for RCPG expression (Fig. 1I).

To verify that BRN2 is responsible for XGA synthesis, we transformed *brn1/2* plants with constructs containing *BRN1-GFP* or *BRN2-GFP*. BRN1-GFP failed to switch on XGA synthesis in *brn1/2* but it activated RCPG expression, rescuing the phenotype of extra BLCs (Fig. 5 K and P). In contrast, XGA was observed in the root cap of *brn1/2* plants transformed with the *BRN2-GFP* construct. *RCPG* was expressed and excess BLCs did not accumulate in *brn1/2/BRN2-GFP* root cap samples (Fig. 5 L and P). These results demonstrate that BRN2 is required for XGA synthesis and both BRN1 and BRN2 can activate *RCPG* expression.

*brn2, and brn1/2/BRN1-GFP* did not exhibit defects in BLC shedding despite that XGA synthesis is blocked in the mutant root cap and that RCPG is a cargo protein of XGA vesicles (Fig. 5, I and K). This observation implied that RCPG secretion remained unaffected in the absence of functional *BRN2*. We investigated the localization of RCPG in *brn2* root cap cells by conducting immunogold labeling with the RCPG antibody. RCPG-specific antibodies were associated with the Golgi cisternae and TGN compartments in the Golgi stacks of brn2 BLCs (Fig. 3 K-N). These findings indicate that *BRN2* is essential for XGA synthesis but suggest that RCPG could be secreted from BLCs through conventional exocytosis from the TGN in *brn2* BLCs, where XGA synthesis is inhibited.

### *BRN2*-mediated Golgi remodeling was required for the overexpression of RCPG-GFP

Since *BRN2* is required for the production of enlarged vesicles carrying XGA, RCPG, and RCPG-GFP, we examined whether *BRN2* is required for RCPG-RFP overexpression. *oeRCPG-GFP* were crossed with *brn1* or *brn2* and their BLCs were stained for XGA epitopes (Fig. 6A-D). XGA synthesis, Golgi hypertrophy, and RCPG-GFP overexpression were not affected at 2 DAG and 6 DAG in *brn1/oeRCPG-GFP* (Fig. 6 A and C). By contrast, all XGA synthesis, Golgi remodeling, and RCPG-GFP overexpression were shut down in *brn2/oeRCPG-GFP* (Fig. 6 B and D). Golgi stacks in BLCs of *brn2/oeRCPG-GFP* resembled those in *brn2* BLCs, lacking the swollen cisternal margins (Fig. 3J).

**Figure 6.**
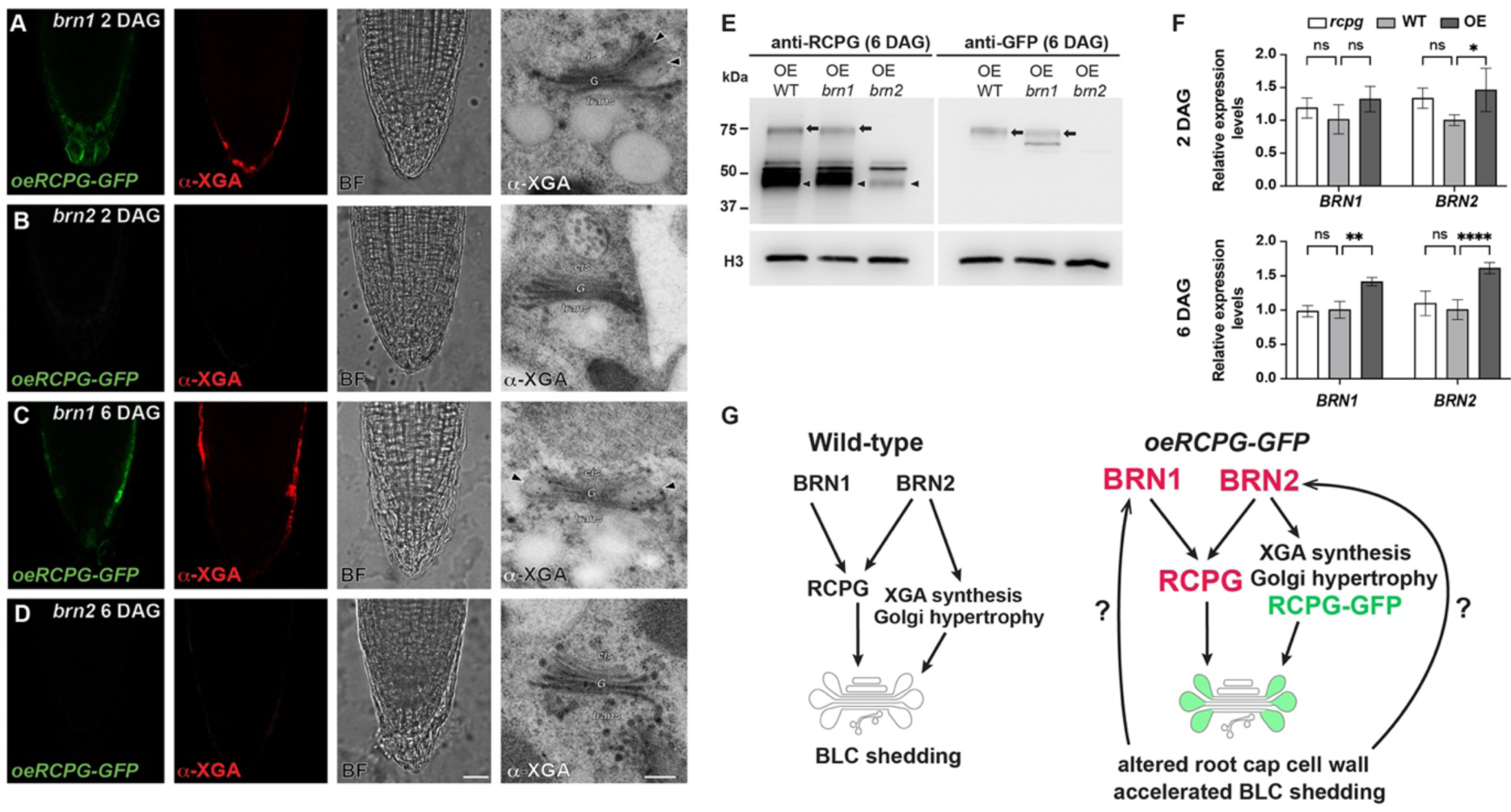
XGA synthesis at 2 DAG and overaccumulation of endogenous RCPG in *oeRCPG-GFP* are dependent on *BRN2*. **A-D.** Whole mount immunofluoresence and immunogold localization of XGA in *brn1/oeRCPG-GFP* (A and C) and *brn2/oeRCPG-GFP* (B and D). GFP, XGA epitopes and XGA-carrying vesicles are not observed in *brn2/oeRCPG-GFP* at 2 and 6 DAG (B and D). The effects of RCPG-GFP overexpression were not affected in the *brn1* background (A and C). Arrowheads indicate XGA-carrying vesicles. Bars in D = 25 μm and 200 nm. **E.** Immunoblot analysis of *oeRCPG-GFP*, *brn1/oeRCPG-GFP* and *brn2/oeRCPG-GFP* with the RCPG or GFP antibody. Arrows and arrowheads indicate RCPG-GFP and endogenous RCPG, respectively. RCPG-GFP was missing and endogenous RCPG did not over-accumulate in *brn2*. **F.** Relative expression levels of *BRN1* and *BRN2* in *rcpg*, wild-type (WT) and *oeRCPG-GFP* (OE) at 2 DAG and 6 DAG. The expression levels were normalized with the levels of GAPDH and analyzed by TWO-WAY ANOVA (ns, p > 0.05. *, p ≤ 0.05. **, p ≤ 0.01. ****, p ≤ 0.0001.). G, A diagram illustrating the BRN2-mediated XGA synthesis and Golgi hypertrophy in Arabidopsis BLCs.

RCPG-GFP polypeptides were detected in the immunoblot analyses of protein samples from *brn1/oeRCPG* and *oeRCPG* but not from *brn2/oeRCPG* (arrows in Fig. 6E). RCPG concentration was higher in *brn1/oeRCPG* than *brn2/oeRCPG* as predicted from excess accumulation of RCPG in *oeRCPG-GFP* (arrowheads in Fig. 6E). These results showed that *BRN2* is essential for the overexpression of RCPG-GFP and upregulation of endogenous RCPG in *oeRCPG-GFP*

Transcript levels of *BRN1* and *BRN2* were compared in *rcpg*, wild-type, and *oeRCPG-GFP* with qRT-PCR (Fig. 6F). mRNA from *BRN2* and *BRN1* were more abundant in *oeRCPG-GFP* at 6 DAG but only *BRN2* exhibited upregulation at 2 DAG, showing that overexpression of RCPG-GFP triggers XGA synthesis and Golgi remodeling by activating *BRN2*.

## Discussion

Our study identifies *BRN2* as a key mediator of the co-secretion of RCPG and XGA. We found *BRN2* was necessary for XGA production, while either *BRN1* or *BRN2* was sufficient for the expression of RCPG as observed in the *brn1* and *brn2* single mutant lines (Fig. 6G). The results from correlative light and electron microscopy and immunogold labeling indicated that RCPG is secreted in conjunction with XGA. Notably, overexpression of RCPG-GFP led to structural alterations in the BLC cell wall, accelerated BLC turnover, and upregulation of *BRN1*, *BRN2*, and endogenous *RCPG*. The activation of *BRN1* and *BRN2* by RCPG-GFP overexpression suggests a positive feedback cycle that facilitates BLC detachment, which is the irreversible terminal stage of the root cap cell maturation. Positive autoregulatory loops are known to play a role in one-way developmental processes in plants, such as leaf senescence and juvenile-to-adult transition (Zhuo et al., 2020; Meng et al., 2021). The self-reinforcement was hindered in *brn2/oeRCPG-GFP*, implying that *BRN2* is crucial for accommodating the elevated levels of RCPG, perhaps through XGA synthesis and Golgi hypertrophy. However, the signaling mechanism underlying the synergistic expression of *RCPG* and *BRN2* remains to be determined. Considering that the truncated RCPG-GFP did not cause surplus RCPG (Fig. S3), export of active RCPG appears to be critical.

A constitutive promoter, *Ubq10*, was adopted for ectopic expression of RCPG-GFP in *oeRCPG-GFP*. Interestingly, *RCPG-GFP* was expressed in BLCs and cells immediately inside BLCs despite its Ubq10 promoter that is active in non-root cap cells (Grefen et al., 2010). The RCPG-GFR positive cells of *oeRCGP-GFP* coincide with the cell type where *BRN2* is expressed (Kamiya et al. 2016). The overlap of RCPG-GFP and *BRN2* promoter activity in the *oeRCGP-GFP* line agrees with the requirement of *BRN2* for the overaccumulation of endogenous RCPG. Furhter investigation is needed to characterize how *BRN2* enables the excess accumulation of RCPG.

### Function of XGA in BLC shedding

We found that RCPG was secreted from the BLCs of *brn2* through the TGN and that BLC separation was not inhibited. In the *brn1/2* double mutant, where RCPG is not expressed, additional BLC layers attached to the root cap were observed, proving that RCPG is the primary factor for BLC shedding. The role of XGA in BLCs remains unclear, although XGA synthesis is coupled to cell wall decay and cell separation in plants (Willats et al., 2004; Mravec et al., 2017; Wang et al., 2017; Wang and Kang, 2018).

One possibility is that XGA could facilitate isolating RCPG from other secretory proteins and packaging RCPG into the enlarged XGA-carrying vesicles in the trans-Golgi. Secretory vesicles produced from the TGN contain regular non-cellulosic polysaccharides and arabinogalactans and they are constitutively secreted to the cell wall (Kang et al., 2011; Kang et al., 2022). Export of RCPG via such vesicles could be detrimental for cell wall maintenance. In agreement with the notion, RCPG epitopes that concentrate to XGA-carrying vesicles relocated to the TGN in *brn2* BLCs where Golgi stacks are devoid of XGA. XGA might have affinity for RCPG but not be hydrolyzed as it has polygalacturonic acid backbone with unique substitutions.

It is worth noting that XGA accumulates in Golgi stacks of all BLC cell walls, while RCPG expression is limited to BLCs in the CRC. This suggests that XGA may play roles not linked to RCPG. XGA also accumulates in border cells of maize, a monocot plant whose primary cell wall has a distinct composition from dicot plants such as Arabidopsis.

### Inhibited Golgi swelling in *brn2*

Our findings suggest that BRN2 is a transcription factor essential for orchestrating the release of BLCs. It activates RCPG and triggers the machinery for XGA synthesis. Root cap maturation is aberrant in *smb/brn1/brn2* triple mutant root cap and Bennett et al. (2010) examined expression of candidate effectors of the NAC-type transcription factors. An Arabidopsis cellulase gene, *CEL5*, was one of the candidates as it is highly transcribed in BLCs of CRC like RCPG. Its expression was suppressed *smb/brn1/brn2* more than *smb*, providing evidence that *CEL5* is a downstream gene of BRN1 and BRN2 (Bennett et al., 2010).

A Golgi-localized aminophospholipid translocase *ALA3* is expressed in the root cap and *ala3* mutant exhibited defects in BLC separation. Additionally, hypertrophied Golgi stacks disappeared in *ala3* root cap cells, as in *brn2*, suggesting that lipid asymmetry is implicated in the swelling of cisternal margin and production of XGA-carrying vesicles (Poulsen et al., 2008). This study implies that downstream effectors of BRN2 may include regulators of vesicular trafficking to accommodate the enhanced secretion from border cells/BLCs. Interestingly, the overexpression of RCPG-GFP was inhibited in the *brn2* mutant, which lacks enlarged Golgi stacks in the root cap.

### Accelerated BLC turnover by RCPG-GFP overexpression

The detachment of BLCs is similar to cell separation that occurs in abscission zones, but in the root cap, the structure must be retained for the continuation of its function after cell loss (Kumpf and Nowack, 2015). This requires communication between maturing BLCs in the distal root cap and the dividing cells in the root cap meristem. In addition, cell division, maturation, and separation should be sequentially coordinated along the proximodistal axis of the root cap (Iijima et al., 2008).

Cell-to-cell signaling involving a small peptide, IDA-like1 (IDL1), has been shown to play a role in BLC renewal (Shi et al., 2018). The small peptide INFLORESCENCE DEFICIENT IN ABSCISSION (IDA) and two closely related leucine-rich repeat receptor-like kinases (LRR-RLKs), HAESA (HAE), and HAESA-LIKE2 (HSL2), control cell separation for floral organ abscission and lateral root emergence (Butenko et al., 2003; Cho et al., 2008; Stenvik et al., 2008; Zhu et al., 2019). In the Arabidopsis root cap, upregulation of IDL1 results in accelerated BLC loss and cell division in the root cap meristem, and the acceleration of BLC turnover requires HSL2 (Shi et al., 2018). RCPG-GFP overexpression resulted in significantly faster BLC turnover than the wild type, but its root cap architecture was not affected. Although RCPG was limited to BLCs of the CRC in wide type, RCPG-GFP accumulated in extra cell layers inside the BLCs and BLCs derived from LRC in *oeRCPG-GFP*. A similar expanded expression was observed for BRN2 and RCPG when IDL1-HSL2 signaling was enhanced in the root cap (Shi et al., 2018). Therefore, RCPG-GFP overexpression is likely to act through the IDL1-HSL2 pathway. An auxin gradient from the proximal to the distal axis of the root cap is also involved in the regulation of cell division and detachment (Dubreuil et al., 2018). Further study of *oeRCPG-GFP* could reveal novel aspects of the control of root cap homeostasis against the cell maturation flux.

## Materials and methods

### Plant materials, growth conditions, genotyping and transformation

The *pRCPG:RCPG-RFP* transgenic line, *rcpg* (GABI_100C05), and *brn1/2* (*brn1-1*, SALK_151986, and *brn2-1*, SALK_151604) were kindly provided by Dr. Tatsuaki Goh (Kamiya et al. 2016). The *brn1-1* (SALK_151986C) and *brn2-1* (SALK_151604C) were ordered from the *Arabidopsis* Biological Resource Center (ABRC). The transgenic lines were screened on the half Murashige and Skoog (MS) plates with 10% sucrose containing 50 mg/L kanamycin. *pUBQ10:RCPG-GFP* was crossed with *rcpg*, *brn1-1* and *brn2-1*, and the second generation seedlings were put into soil for the DNA extraction with Edward buffer (Edwards et al., 1991). The homozygous mutants were identified by genotyping with the primers listed in Table S1 using Vazyme 2x Taq Master Mix on Bio-Rad C1000 Touch^TM^ Thermal Cyclers.

All *Arabidopsis* seeds were sterilized with 75% ethanol for 5 minutes twice, and 100% ethanol for 1 min. The seeds were then washed with sterile distilled water three times and cultured on half MS agar plates without sucrose. The plates were grown vertically in a growth chamber with a light intensity was 120-150 µmol/m^2^ at 24°C, under a 16-hour light and 8-hour dark cycle.

The agrobacterium strain GV3101 was used for plant transformation, and the plasmids were transformed into agrobacterium by electroporation via Bio-Rad Gene Pulser Xcell Electroporation Systems. To create transgenic plant lines, we peformed agrobacterium-mediated floral dip method (Zhang et al. 2006).

### RNA extraction, cDNA synthesis, cloning and RT-qPCR

Root tip samples (∼80) were harvested for RNA extraction by TRIzol^TM^. cDNA was reverse transcribed from 1μg of total RNA by Bio-Rad iScript^TM^ gDNA Clear cDNA Synthesis Kit. The transgenic lines were generated using binary vector *pBI121-GFP*. The original *CaMV 35S* promoter on the vector was replaced with *ubiquitin10 (UBQ10)* promoter for RCPG constitutive expression. The coding DNA sequence (CDS) of RCPG, BRN1, and BRN2 were amplified from Arabidopsis cDNA, and truncated RCPGs were generated by overlapping PCR from RCPG CDS fragment. RCPG (ΔC) (1-57 aa) excluded the GH28 domain (58-397 aa) of RCPG, while RCPG (ΔL) (1-20 + 58-397 aa) combined the signal peptide (1-20 aa) with the GH28 domain. The fragments were separately inserted into the same position between the restriction enzyme sites of BamH I and Kpn I (NEB) on the reconstructed *pUBQ10:GFP pBI121* or original 35S:GFP pBI121 vector at the N-terminal of GFP based on a previous study (Kamiya et al. 2016). SsoAdvanced Universal SYBR Green Supermix was used in the Reverse Transcription Quantitative PCR (RT-qPCR), and Ct (cycle threshold) values were measured by Bio-Rad CFX96 Touch Real-Time PCR Detection System. ΔΔCt method was performed for calculating relative gene expression level, and one/two-way ANOVA were used in differential analysis. *GAPDH* was used as reference gene (Dekkers et al., 2012), and the primers used are listed in Table S1.

### Cryopreservation, TEM, and electron tomography

*Arabidopsis* seeds were grown vertically on half MS agar plates without sucrose. The root tips were dissected and transferred into 3 mm planchettes with 0.15 M sucrose as the cryoprotectant. The samples were frozen using High Pressure Freezer Leica EM ICE followed by freeze-substitution at −80 °C for 24-48 hours in Leica EM AFS2 machines. For ultrastructural analysis, samples were fixed in in 2 % osmium tetroxide (OsO4) dissolved in acetone, while for immunogold labeling and immunofluorescence experiments, samples were processed in acetone containing 0.25 % glutaraldehyde and 0.1% uranyl acetate.

For ultrastructural analysis, the samples were first kept at 4 °C overnight, then transferred to room temperature for one hour and washed with acetone every 15-30 minutes for three times. They were then embedded in Epon resin with a gradient dilution in acetone (10, 25, 50, 75 and 100%) over three days and polymerized at 60°C overnight. For immuno-samples, the samples were embedded in Lowicryl HM20 resin, and polymerized under an ultraviolet lamp overnight in the AFS machines at −45°C after being washed three times with acetone and substituted in an increased concentration of HM20 resin (33, 66, and 100%, diluted in acetone) over three days (Kang, 2010). TEM and electron tomography were carried out with Hitachi H-7650 TEM (Hitachi-High Technologies, Japan) operated at 80 kV and Tecnai F20 electron microscope (Thermo-Fischer, USA) operated at 200 kV, respectively as described in (Liang et al., 2022) and (Mai et al., 2019).

### Immunogold labeling and CLEM

Thin sections (90-200nm) of HM20 samples were cut using the Leica EM UC7 Ultramicrotome and collected on nickel slot grids coated with 0.75% formvar. The grids were floated on 0.1N HCl drops to remove the glutaraldehyde and increase the specificity of labeling in the humid chamber. Subsequently, the grids were incubated in 2% Bovine serum albumin (BSA) dissolved in either PBS or PBST (PBS for immunofluorescence and PBST for immunogold labeling) for 30 minutes. After blocking, the grids were probed with primary antibodies diluted in 1% BSA overnight at 4°C. The probed grids were washed by 0.5% BSA three times and then incubated with gold-particles/fluorescent-dye conjugated secondary antibodies, which were diluted in 0.5% BSA for one hour. Following this, the grids were washed with PBS/PBST to remove the non-specific binding. For CLEM samples, the washed grids were kept on the PBS drops in the dark humid chamber at 4°C until they were ready for imaging using the confocal microscope (Kang, 2010; Wang et al., 2019).

For double-immunogold-labeling/immunofluorescence, the XGA antibody (LM8) was mixed with anti-GFP/RFP, or α-XG. The dilutions of primary antibodies were 1:10 for GFP and RFP antibodies, 1:15 for LM8, 1:20 for LM1, and 1:20 for α-XG. The dilution of secondary antibodies was 1:10.

### Whole-mount Immunofluorescence and time-lapse imaging

The whole-mount immunofluorescence was performed as previously described (Sauer et al., 2006). *Arabidopsis* seedlings at 2 DAG and 6 DAG were fixed with 4% paraformaldehyde at room temperature and probed with LM8 overnight at 4°C. After washing with PBS, the samples were incubated with secondary antibody at 37°C for three hours. The dilution of LM8 was 1:150, and the dilution of the secondary antibody was 1:100.

### Root cap cuticle staining

The root cap cuticle (RCC) of 8-19 seedlings were firstly stained with Fluorol Yellow (FY) 088 (0.01% in methanol) at 65 °C for 30 minutes, followed by washing with distilled water. Then the seedlings were counterstained with aniline blue (0.5% in water) in dark condition at room temperature for one hour. After the counterstaining, seedlings were washed in distilled water and transferred to confocal microscope for observation (Naseer et al., 2012; Berhin et al., 2019).

### Propidium iodide (PI) staining, and cell length analysis

2 DAG and 6 DAG seedlings were stained with PI for 5-10 minutes and washed with distilled water before went to microscope. The PI solution was diluted in distilled water for the 10 ug/mL working concentration. Z-stack series images near the central plane of root tips were captured by Leica SP8, and the center-most images were chosen for calculating the length of root cap cells. Around 20 serial images were taken for each root, and 10-16 roots were stained with PI for each group. The cell length was measured by ImageJ, and the differential analysis was performed using two-way ANOVA.

### BFA treatment and FM4-64 staining

BFA treatment was performed using 25μM Brefeldin A (BFA) and 5μg/mL FM4-64 as described previously (Uemura et al., 2012). *Arabidopsis* seedlings were incubated in 60 mm petri dish with 25 μM BFA diluted in half MS liquid medium for 2 hours. Then the seedlings were transferred to FM4-64 containing petri dish for 15 minutes and washed with distilled water before imaging.

## Supplemental data

**Figure S1.**
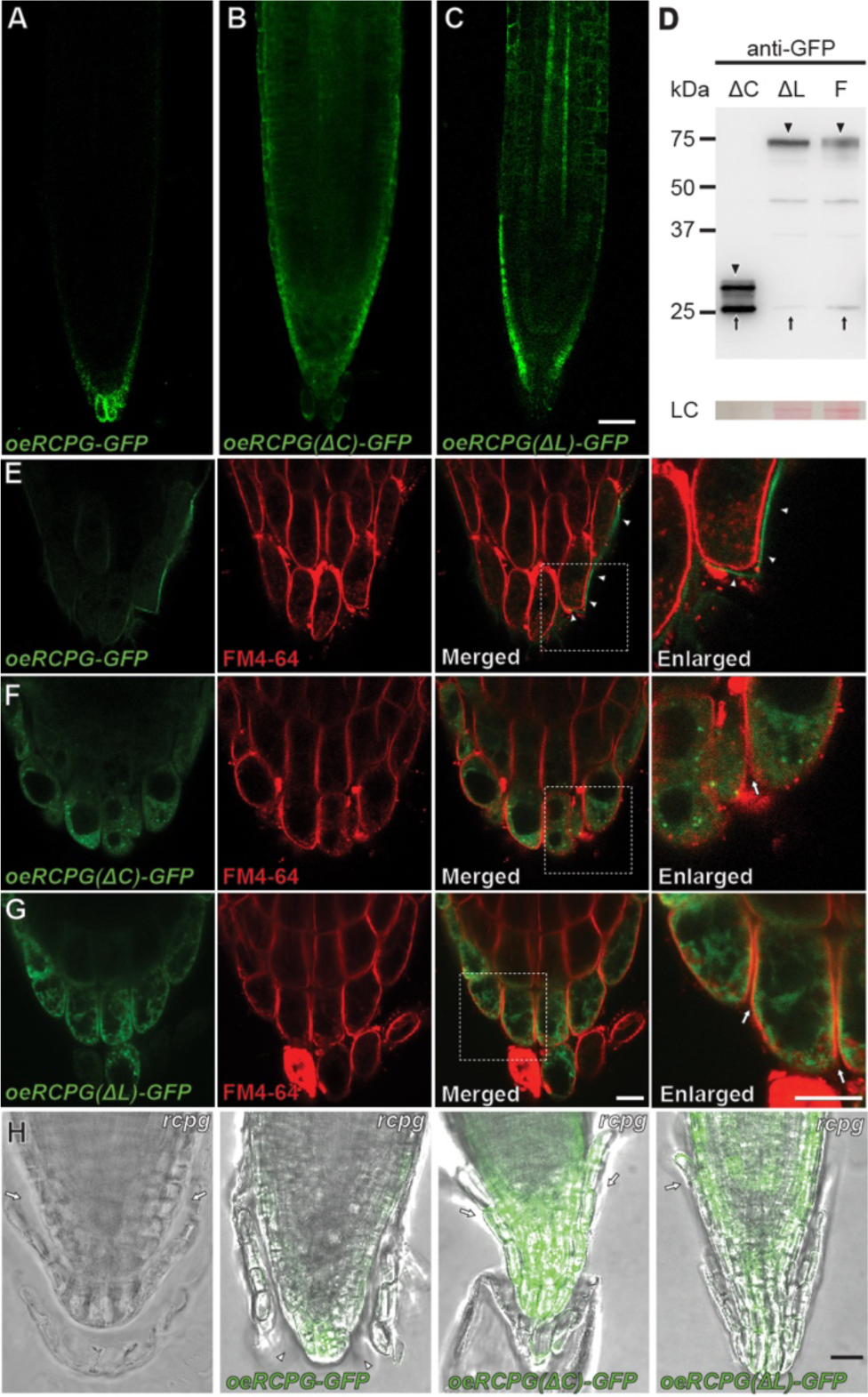
Truncated forms of RCPG were not exported to the cell wall and they failed to rescue the *rcpg* mutant phenotypes. **A-C.** Root tips of transgenic lines expressing RCPG-GFP, RCPG(ΔC)-GFP (lacking the catalytic domain) RCPG(ΔL)-GFP (lacking the linker region) with the *UBQ10* promoter. Full length RCPG-GFP fusion proteins are seen in BLCs (*oeRCPG-GFP*), while truncated RCPG-GFP versions accumulate in the root cap as well as root meristem cells. Bar = 25 μm. **D.** immunoblot with an anti-GFP for detecting GFP fusion proteins (arrowheads) and free GFP (arrow). Ponceau S staining is shown as the loading control (LC). **E-G.** Root caps of three transgenic lines after FM4-64 staining. RCPG-GFP is exported to the cell wall (*i.e.*, outside the plasma membrane stained with FM4-64, arrowheads in E). Truncated RCPG-GFPs are retained in BLCs. (F and G, arrows). oeRCPG(ΔC)-GFP exhibits cytoplasmic puncta distinct from the FM4-64 stained endosomal compartments. oeRCPG(ΔL)-GFP exhibits a network-like fluorescence, typical of the endoplasmic reticulum. Bars = 10 μm. **H.** *rcpg* expressing full-length RCPG-GFP, RCPG(ΔC)-GFP, or RCPG(ΔL)-GFP. The full-length RCPG-GFP rescued the *rcpg* phenotype of extra BLC layers (arrowheads), but the truncated RCPG-GFPs did not (arrows). Bar = 25 μm.

**Figure S2.**
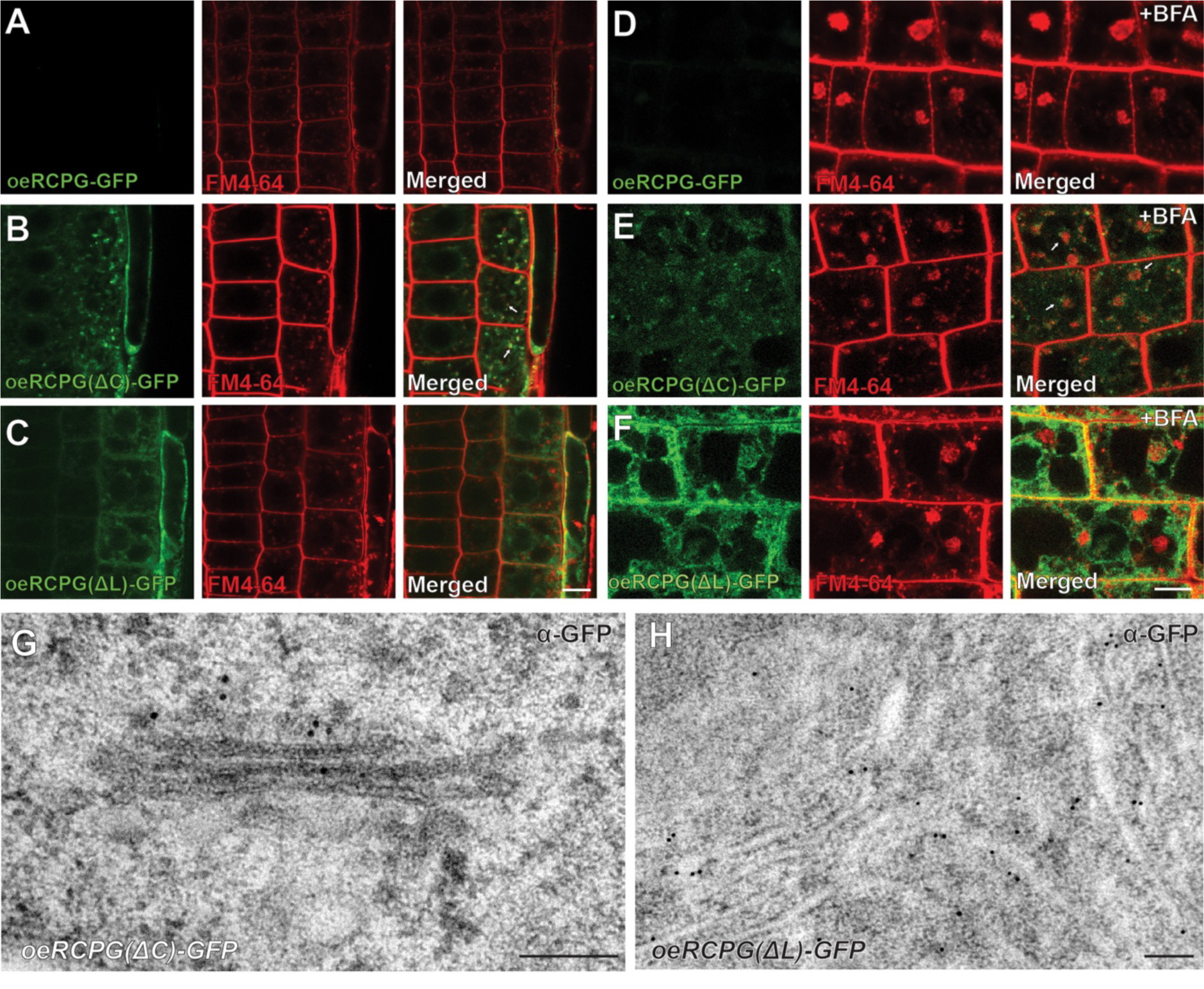
Truncated forms of RCPG were retained in the *cis*-Golg or the ER. **A-C.** Root meristem cells of transgenic lines expressing *RCPG-GFP*, RCPG-GFP without the catalytic domain (1′C), RCPG-GFP without the linker domain (1′L) with the *UBQ10* promoter. GFP fluorescence was not detected in meristem cells expressing the full length RCPG-GFP despite that the fusion protein synthesis was driven by the Ubq10 promoter (A). RCPG(1′C)-GFP and RCPG(1′L)-GFP were associated with cytosolic puncta (arrows in B) and network, respectively (B and C). **D-F.** The three transgenic lines after brefeldin A (BFA) treatment. The oeRCPG(ΔC)-GFP puncta surrounded BFA bodies (E, arrows). Bars = 7.5 μm. **G-H.** Immunogold labeling the *oeRCPG(ΔC)-GFP* (G) and *oeRCPG(ΔL)-GFP* (H) lines. oeRCPG(ΔC)-GFP and oeRCPG(ΔL)-GFP gold particles localized to the *cis*-Golgi (G) and ER (H), respectively. Bars=200 nm.

**Figure S3.**
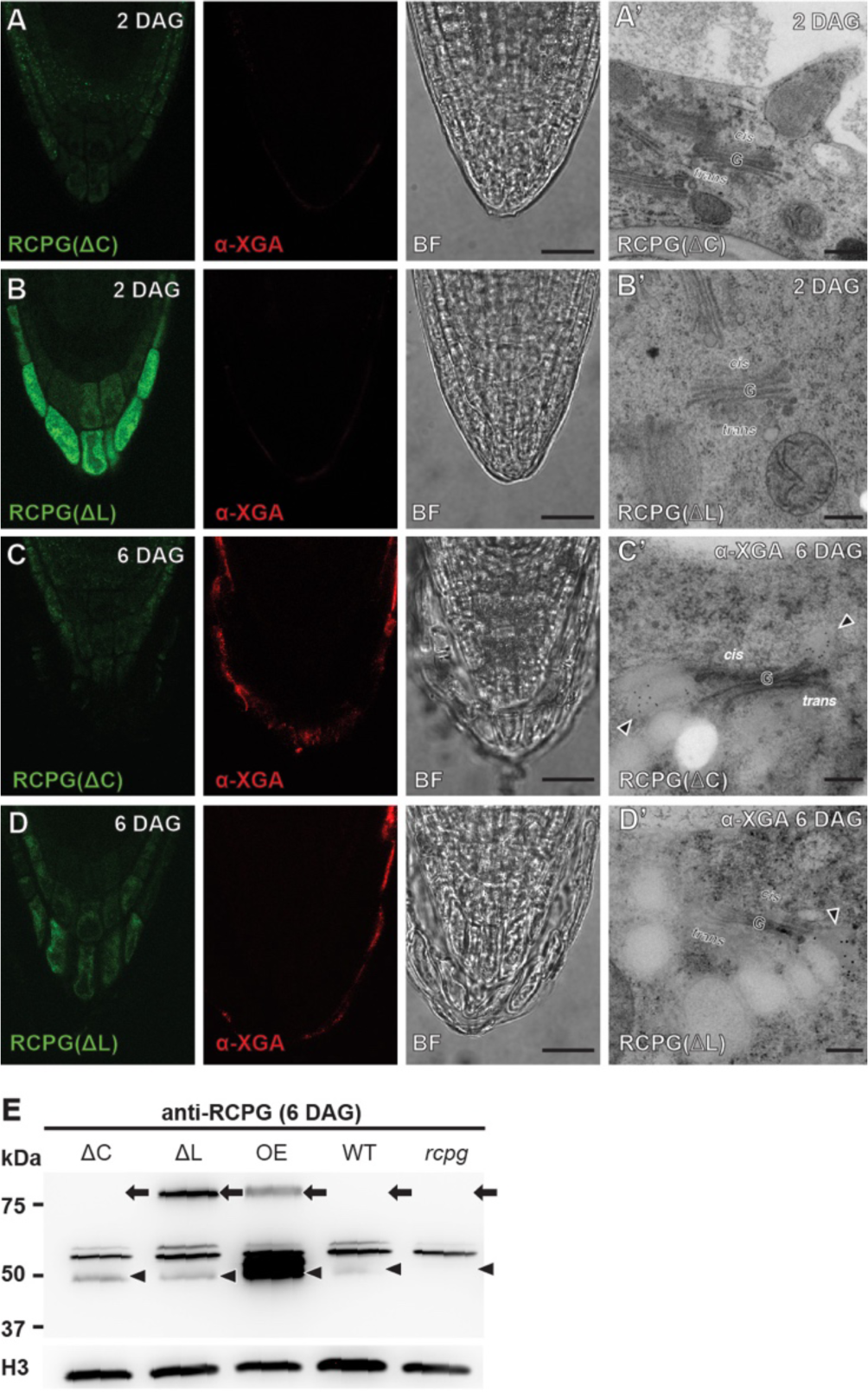
Overexpression of RCPG(ΔC)-GFP and oeRCPG(ΔL)-GFP did not induce ectopic production of XGA-carrying vesicles or overaccumulation of RCPG. **A-D.** Whole mount immunofluorescence localization of XGA with LM8 (XGA-specific antibody). Root tip samples from *oeRCPG(ΔC)-GFP* and *oeRCPG(ΔL)-GFP* were examined at 2 DAG and 6 DAG. XGA epitopes were not noticed in the transgenic lines (A-B) and they did not have hypertrophied Golgi stacks (A’-B’) at 2 DAG. Bars = 50 μm. Bars in A’ to D’ (electron micrographs) = 200 nm. **E.** Immunoblot analysis of RCPG-GFP lines with the RCPG antibody. No excess endogenous RCPG was observed in *oeRCPG(ΔC)-GFP* or *oeRCPG(ΔL)-GFP*. Arrowheads and arrows mark sizes of native RCPG and RCPG-GFP, respectively. Histone H3 was used as the loading control.

**Supplemental Table S1.** Primer sequences

**Supplemental video S1**. Time-lapse movie of Arabidopsis root tip expressing RCPG-GFP with the *RCPG* native promoter.

**Supplemental video S2.** Time-lapse movie of Arabidopsis root tip expressing RCPG-GFP with the *UBQ10* promoter.

